# Temporal Pattern of Mutation Accumulation in SARS-CoV-2 Proteins: Insights from Whole Genome Sequences Pan-India Using Data Mining Approach

**DOI:** 10.1101/2023.07.07.548087

**Authors:** Chakrakodi N Varun

## Abstract

Mutation is a fundamental factor that affects host-pathogen biology and consequently viral survival and spread. Close monitoring and observation of such mutation help decipher essential changes in the SARS Cov2 genome. A plethora of mutations have been documented owing to increased whole genomic sequencing. Understanding how conserved the specific mutations are and the temporal pattern of mutation accumulation is of paramount interest. Using an in-house data mining approach, pan-India data was mined and analysed for 26 proteins expressed by SARS-CoV-2 to understand the spread of mutations over 28 months (January 2021-April 2023). It was observed that proteins such as Nsp3, Nsp4, ORF9b, among others, acquired mutations over the period. In contrast, proteins such as Nsp6-10 were highly stable, with no detectable conserved mutations. Further, it was observed that many of the mutations that were highly prevalent in the delta variants were not observed in the omicron variants, which probably influenced the host-pathogen relationship. The study attempts to catalogue and focus on well-conserved mutations across all the SARS-CoV-2 proteins, highlighting the importance of understanding non-spike mutations.

## 1. Introduction

Mutation and evolution are fundamental processes that influence host-pathogen biology and, thereby, virus survival and spread. Constant surveillance and monitoring to decipher new and high-frequency mutations in the SARS-CoV-2 genome are thus essential to understand their adaptation. SARS-CoV-2 which was previously declared as a public health emergency (1), has recently been de-escalated from its pandemic status by the WHO (2). Understanding the dynamics of these mutations over a more extended period is essential and provides significant leads to understanding virus evolution. With the availability of a rapidly increasing number of open-access whole genome sequence data for SARS-CoV-2, there has been a growing interest in data mining and exploring genetic variations (3,4).

The 30 Kbp SARS-CoV-2 genome consists of a single-stranded + sense RNA. The genome encodes 12 open reading frames (ORFs), supporting the expression of 22 non-structural proteins (NSPs) and four structural proteins (spike, envelope, membrane and nucleocapsid). Further, novel coding transcripts have also been proposed (5–7). The SARS-CoV-2 mutation rate is estimated at 10^−6^ mutations per nucleotide per replication cycle (8). However, various parts of the genome accumulate mutations at various degrees, probably due to selection pressure. For example, spike protein, the primary target of vaccines, has been rapidly evolving over the period.

Though a vast volume of published literature catalogues various mutations in various genes of SARS-CoV-2, many of these mutations have occurred in selective subvariants or at low frequencies, thereby losing focus on conserved mutations over a longer period across multiple proteins. The analysis presented herein provides a catalogue of mutations on the SARS-CoV-2 evolution at an amino acid sequence level. In summary, pan-India data was mined and analysed for 26 proteins expressed by SARS-CoV-2 to understand the breadth of conserved mutations over 28 months (January 2021-April 2023). As expected, the spike shows a rapidly accumulating number of mutations; other proteins, such as nsp3 and nsp4, show a slow accumulation of mutations over the period. Parallelly, Nsp’s 7-10 were found to be highly stable.

## 2. Methodology

### 2.1 Data Collection and raw data availability

All the genomics and metadata reported in the analysis herein were downloaded from the GISAID’s EpiCoV database. In brief, the database was searched for SARS-CoV-2 original, high-coverage sequences for samples collected from January 2021 to April 2023. A total of 71,878 sequences were downloaded for screening. The relevant GISAID Identifiers and their doi are provided in Supplementary data 1.

### 2.2 Data pre-processing for downstream analysis

The data were screened manually, and partial genome sequences were removed from further analysis. The whole genome data was then subject to analysis using the Nextclade tool (v2.13.0) (9) using Wuhan-Hu-1/2019 (MN908947) as a reference sequence. Month-wise protein sequences were derived into a multi-fasta file. The overall qc (quality control) status for each sequence, based on the qc score, was obtained in a .tsv file. Only those sequences assessed as “good” by the nextclade algorithm were taken for further downstream analysis. The workflow summary is presented in Figure 1.

**Figure 1:**
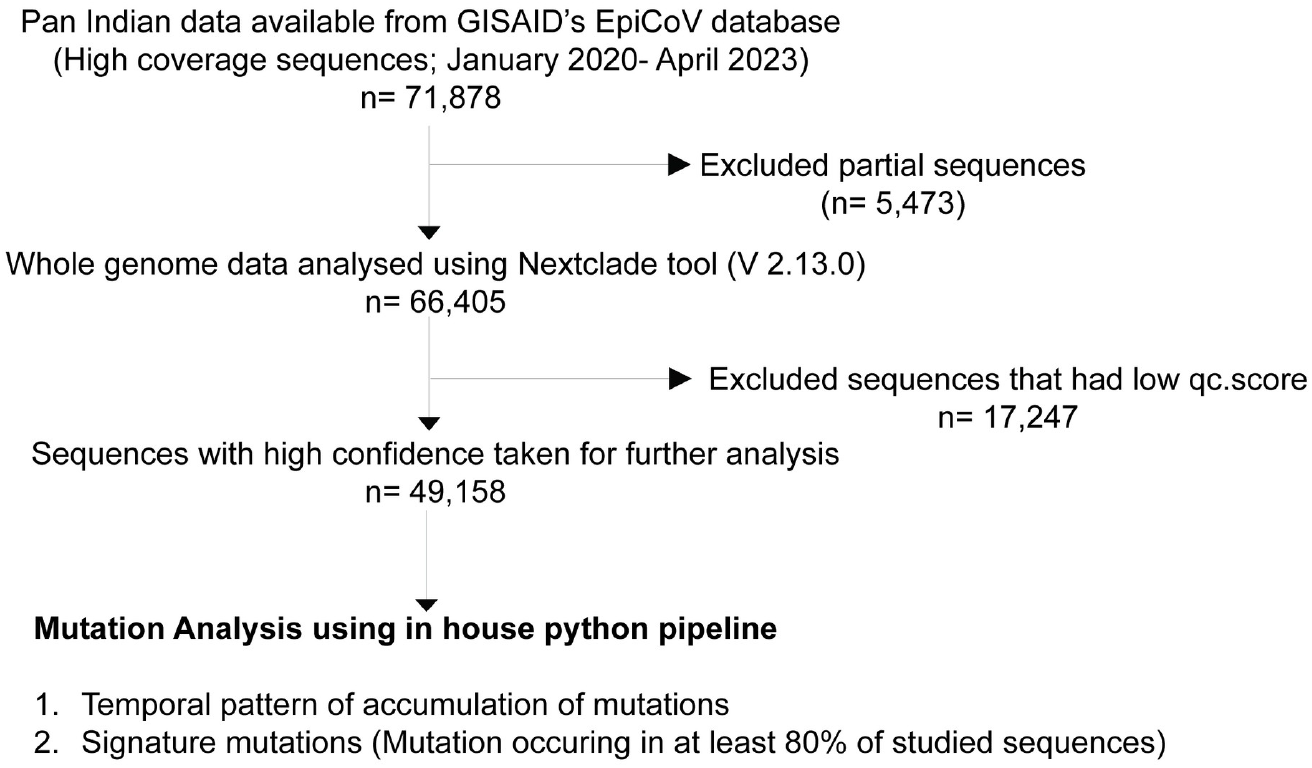
Summary of the data mining and analysis methodology.

### 2.3 Mutation analysis

The aligned protein sequences for all the proteins, month-wise, were analysed using an in-house developed program. In brief, the program was written using Jupyter notebook (v 6.4.12; Anaconda Navigator-2.3.2). The individual sequences were parsed using biopython (v1.81) (10). The codes were further optimised using the “pandas” library. Each sequence that scored as “good” in the nextclade was piped for further analysis. For nsp1-16, the ORF1ab preprotein was computationally spliced into individual proteins based on the published coordinates (11). Nsp11 and ORF8 were ignored for analysis. Nextclade output-derived multi-fasta files were directly used as an input for all the protein mutation analysis. Each amino acid site between the sample and reference sequence was checked for mutation using a mutation function as given below. Wherever the amino acid was unknown (X), the code ignored them as a mutation. The total mutation for each protein sequence was imported into a .xls file.

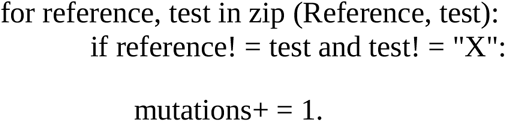

### 2.4 Identification of signature substitution mutations

The pipeline for studying substitution mutations of interest was developed using python and biopython scripts and further optimised using “pandas” library. In essence, the program looked for differences between the reference and the test sequence and reported amino acid change by position, which occurred in at least 80% of the analysed sequences.

## 3. Results & Discussion

Of the total 71,878 sequences that were retrieved-66,405 sequences had a full-length genome sequence suitable for downstream analysis. Amongst these, 49,158 sequences with sample collection dates ranging from January 2021-April 2023 had a good qc status and were taken for further analysis.

### 3.1 S protein

A highly mutating protein generally indicates an active evolutionary process. As already reported by several groups, the S-protein accumulated multiple mutations over the period. (Figure 2A). Since there is no international defining standard cut-off for the frequency of mutations to designate them as “conserved”, I chose 80% as calculative cut-off for determining “highly conserved” mutations. A set of mutations observed in some of the critical lineages reported from India is depicted in Figure 2B. For instance, more recently found strains, XBB 1.16 and XBB 1.15 were found to carry F486P and F490S mutations. F486P enhances SARS-CoV-2 binding to the ACE2 receptor (12). F490S, previously noted in the lambda variant, confers the ability to escape vaccination immunity (13). One of the first mutations, D614G was consistently observed across all the studied sequences (14).

**Figure 2:**
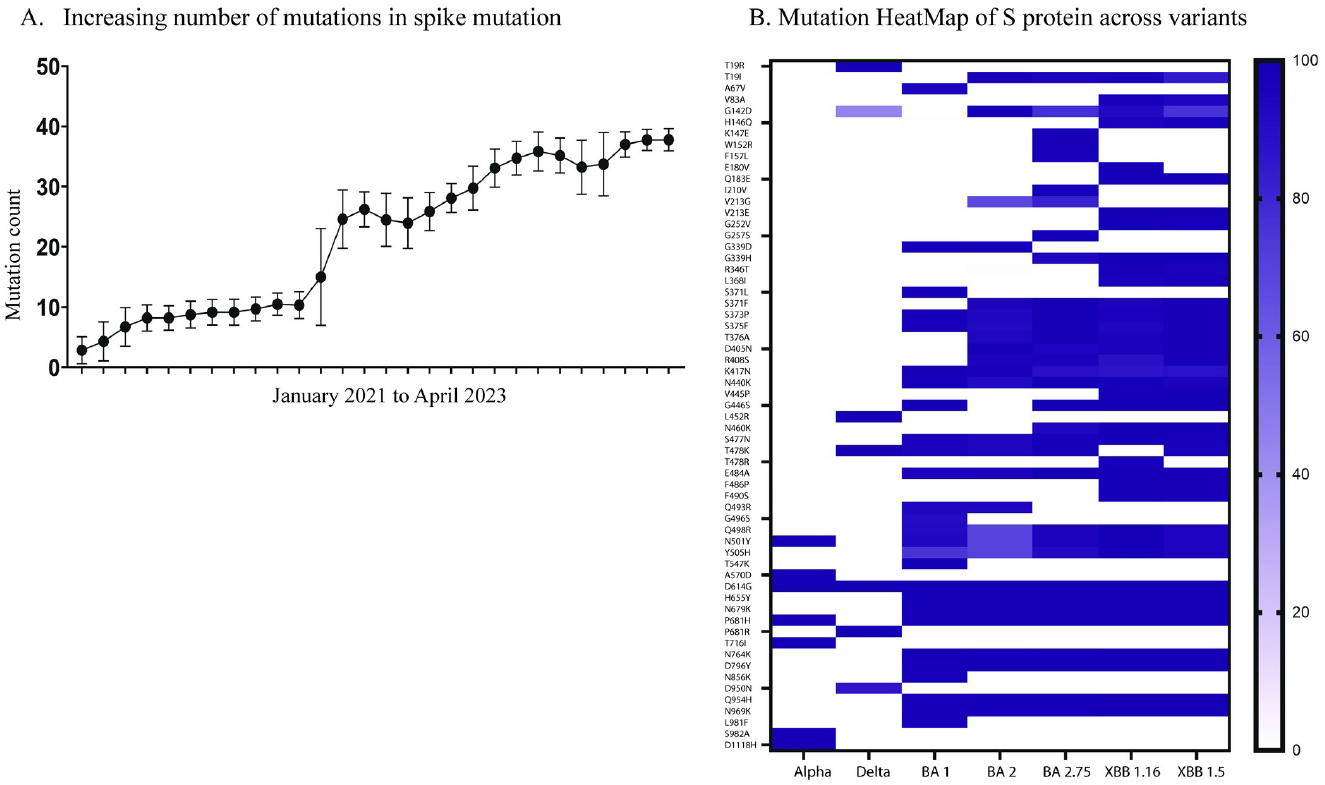
Accumulation of Mutation in SARS-CoV-2 spike protein over the period. **A**. Accumulation of mutations in SARS-CoV-2 spike protein over time. X-axis represents by Month; Y-axis represents the number of mutations. Mean and standard deviation are depicted. **B**. SARS-CoV-2 Spike protein mutations across critical variants. The scale bar represents percentage of sequences showing the mutation.

### 3.2 N, E and M protein

Though there was a rapid increase in the number of mutations in N protein at the initial period, the number of mutations remained stable over a longer time (Figure 3A). However, the specific amino acid mutations differed between delta and omicron variants. The D63G mutation, highly prevalent in delta variants, was not observed in omicron variants. In contrast, the mutation G204R not observed in delta was commonly noted in omicron (Table 1).

**Figure 3:**
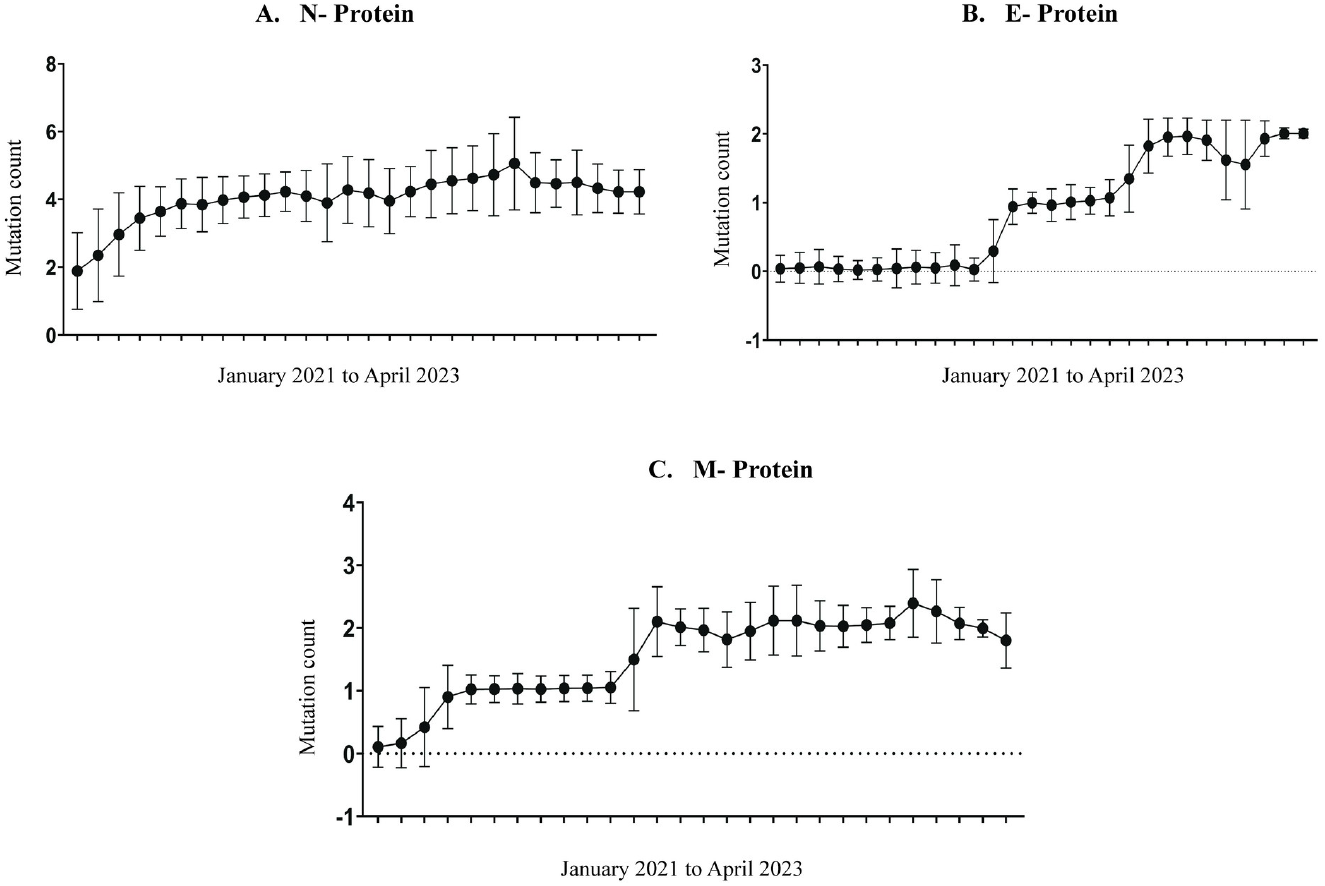
Accumulation of Mutations in N, E and M proteins. Accumulation of mutations in N, E and M proteins (A-C) over time. X-axis represents by Month; Y-axis represents the number of mutations. Mean and standard deviation are depicted.

**Table 1:** Conserved substitution mutations (> 80% sequences carrying the mutation) in N, E and M protein delta, omicron variants and XBB 1.16 subvariant.

| SARS-CoV-2 Protein | Delta and all the subvariants | Omicron and all the subvariants | XBB.1.16 subvariant |
| --- | --- | --- | --- |
| **N protein** | D63G, R203M, D377Y | P13L, R203K, G204R, S413R | P13L, R203K, G204R, S413R |
| **E protein** | - | T9I | T9I, **T11A** |
| **M protein** | I82T | Q19E, A63T | Q19E, A63T |

E protein showed random mutations in the earlier period and later acquired a stable T9I mutation. The T9I mutation is expected to reduce the apoptotic effects on the cell (15). Recently, an additional T11A mutation was observed, which is proposed to reduce cell lethality and lung damage in mouse models (16). M protein showed mutation accumulation similar to E protein. I82T mutation acquired very early, and enhances glucose uptake during viral replication (17), was not observed in omicron. Together, the data suggest that the omicron variant mutations are favoured to be milder than their delta counterpart.

### 3.3 Non-Structural proteins (Nsp’s)

SARS-CoV-2 ORF1ab encodes a total of 16 non-structural proteins. SARS-CoV-2 Nsps are produced biologically by proteolytic cleavage of larger polyproteins. The individual protein sequences were similarly derived using computational slicing as detailed under methodology. Nsp11 encodes a short 13 aa protein, similar to the first segment of Nsp12 and hence was ignored for mutation analysis. It has been previously recognised that ORF1ab accumulated significant mutations, many of which have co-evolved with spike mutations (18,19). It was observed that Nsp1, Nsp3, Nsp4, Nsp5, Nsp12, Nsp13, Nsp14 and Nsp15 accumulated a few significant sets of mutations (Table 2). Other Nsp’s, including Nsp2, Nsp 7-10 and Nsp16, had an extremely low rate of mutation and no recognisable conserved mutations.

**Table 2:** Conserved substitution mutations (> 80% sequences carrying the mutation) in various SARS-CoV-2 Non-structural proteins that were observed in the studied sequences.

| SARS-CoV-2 Protein | Delta and all the subvariants | Omicron and all the subvariants | XBB.1.16 subvariant |
| --- | --- | --- | --- |
| Nsp1 | - | S135R | **K47R**, S135R |
| Nsp3 | - | T24I, G489S | T24I, G489S |
| Nsp4 | - | L264F, T327I, L438F, T492I | L264F, T327I, L438F, T492I |
| Nsp5 | - | P132H | P132H |
| Nsp6 | - | - | L260F |
| Nsp12 | P323L, G671S | P323L | P323L |
| Nsp13 | P77L | R392C | **S36P**, R392C |
| Nsp14 | - | I42V | I42V, **D222Y** |
| Nsp15 | - | T112I | T112I |

**Table 3:**
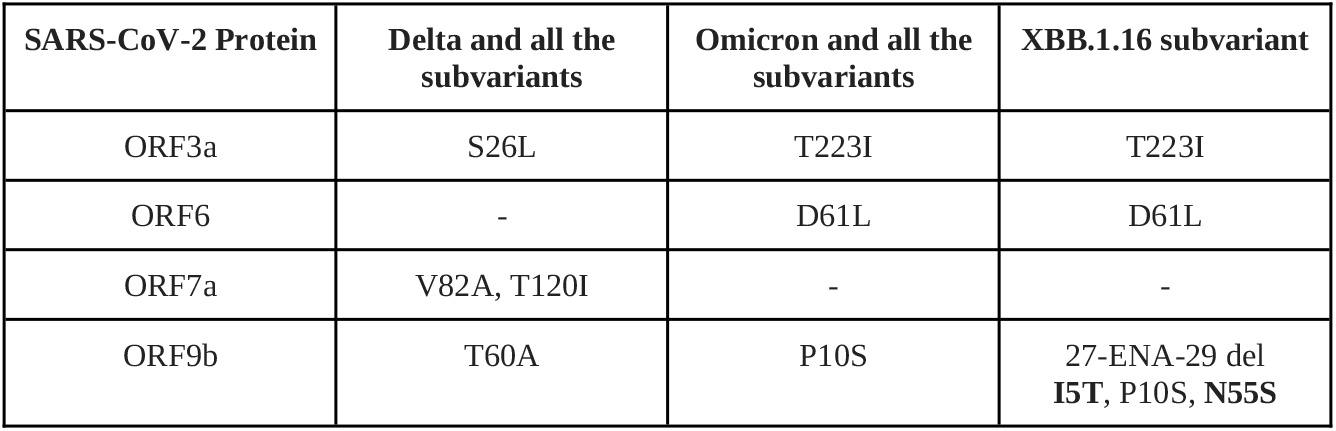
Conserved mutations (> 80% sequences carrying the mutation) in various accessory proteins.

| SARS-CoV-2 Protein | Delta and all the subvariants | Omicron and all the subvariants | XBB.1.16 subvariant |
| --- | --- | --- | --- |
| ORF3a | S26L | T223I | T223I |
| ORF6 | - | D61L | D61L |
| ORF7a | V82A, T120I | - | - |
| ORF9b | T60A | P10S | 27-ENA-29 del **I5T**, P10S, **N55S** |

Notably, the G671S mutation in the RNA-dependent RNA polymerase (RdRp; Nsp12) prevalent in delta is not observed in omicron. G671S mutation facilitates enhanced replication and increased transmission (20). Further, a three amino acid deletion (ΔSGF) mutation in Nsp6 (106-108) was observed in more than 80% of omicron variants. Omicron-derived Nsp6 and spike together have shown a preferential upper respiratory tract infection (21). The evidence of these mutation patterns indicates a propensity of omicron towards causing a milder infection. It should be noted that with new emerging variants, this scenario is changing further. For instance, additional mutations such as K47R (Nsp1), L260F (Nsp6) and S36P (Nsp13) were observed in XBB1.16 for which significance is currently not well understood.

### 3.4 Other accessory proteins

ORF3a, a large accessory protein, over the time period has shown several mutations occurring at a low frequency. Previous studies have suggested several mutations (22), most of which were observed at a low frequency. S26L was found to be a well-conserved mutation earlier, while omicron variants carry T223I, which is predicted to be a destabilising mutation though its functional relevance is unclear (23). ORF6 has acquired a stable mutation-D61L, which probably reduces the ability of the virus to counteract innate immune response (24). ORF7a showed two conserved mutations in delta variants-V82A and T120I. Both these mutations were not present in the omicron variants. Since these mutations occur within the functional domain, the mutational drop is expected to reduce virulence (25). ORF7b has been largely stable, with no identifiable conserved mutations.

ORF9b has recently accumulated a few significant mutations. The accessory protein is known to localise on the mitochondrial membrane and modulate the IFN-I response (26). Currently, the effects of these mutations are unclear. N55S mutation maps to the region that interacts with TOM70, while the I5T mutation is located in the N terminal. Hence, it is speculated that these mutations will modulate the TOM70-ORF9b structure (18).

A detailed mutation analysis for ORF8 protein was not performed since many sequences showed the appearance of stop codons leading to nonsense mutations. It has been suggested that ORF8 is an immunomodulator but not an essential protein (27); hence, loss of protein expression is likely to be of little or no consequence. It was observed that the number of sequences that showed a premature stop codon increased over the period. Indeed, about 91% of the XBB.1.16 sequences showed a stop codon at the 8th position (G8*).

## 4. Limitations

The data mining relies mainly on the open data available through the GISAID database and the nextclade analysis tool. Firstly, the GISAID database probably does not host all the SARS CoV2 whole genome sequences (28,29). Second, the nextclade analysis tool assigns a qc status based on certain assumptions in analysis, which may not always be true (9). Third, the analysis pipeline was designed to omit amino acid sequences designated “X” since they represent an unknown amino acid prediction. Though QC scoring and filtering ensures to remove a majority of those sequences with a large number of “X” in them which may bias the overall analysis, it is acknowledged that those with a good score can still have a few unknown amino acids. Since there is an equal probability that the amino acid is the wild type or a mutation, as a benefit of the doubt, they have not been accounted for in the analysis. However, this represents a fractional number and is thus unlikely to bias the analysis. Further, assuming that the GISAID holds most of the data and nextclade QC scoring helps avoid analysis of sequences with lesser confidence, the limitations does not significantly affect the data presented. Further, as already discussed there are currently no standards available to define a mutation occurrence as “highly conserved”. Since 80% was taken as a cut-off, it is possible that some of the mutations are not captured in the resulting datasets. For example, a spike mutation-T95I, was present in about 74% of the BA.1, but was not captured due to the set cut-off, and hence the data should be interpreted accordingly.

## 5. Conclusion

A massive number of mutations have been reported across various proteins of SARS-CoV-2 thereby loosing focus on which mutations are otherwise important. The strength of the analysis presented herein lies in the breadth of data analysed across the period allowing for the confident identification of significantly conserved mutations. An overall theme that also emerged during the analysis was that various mutations which probably conferred higher virulence to the delta variant were not observed in omicron variants. However, mutations that render higher transmissibility were observed more commonly in omicron variants. It should also be noted that omicron lineage arose independently of delta lineage; hence, they do not likely share many mutations of interest. Additionally, mutations in various other proteins including Nsp1, Nsp13 and ORF9b are seen in recent sequences such as XBB.1.16 which needs to be further evaluated. This analysis highlights the importance of ongoing data mining to observe the acquired and conserved mutations and ignore otherwise seemingly random sets of mutations.

## Acknowledgement

I wish to acknowledge the efforts by the Indian SARS-CoV-2 Genomics consortium (INSACOG) which is largely responsible for SARS-CoV-2 whole genome sequencing in India that has generated the sequences available at GISAID database.

