## Supplementary data 1 for "Temporal Pattern of Mutation Accumulation in SARS-CoV-2 Proteins: Insights from Whole Genome Sequences Pan-India Using Data Mining Approach"

**GISAID Identifiers for data used in this study**

| **Time Period** | **EpiSet** | **doi** | **Number of sequences** |
| --- | --- | --- | --- |
| January 2021- December 2021 | EPI_SET_230327rv | 10.55876/gis8.230327rv | 54,664 |
| January 2022- December 2022 | EPI_SET_230401wt | 10.55876/gis8.230401wt | 15609 |
| January 2023 | EPI_SET_230522co | 10.55876/gis8.230522co | 71 |
| February 2023 | EPI_SET_230403tx | 10.55876/gis8.230403tx | 145 |
| March 2023 | EPI_SET_230424be | 10.55876/gis8.230424be | 938 |
| April 2023 | EPI_SET_230522fe | 10.55876/gis8.230522fe | 451 |
